# The Vaginal Microbiome in Women of Reproductive Age with Healthy Weight versus Overweight/Obesity

**DOI:** 10.1101/2021.03.18.435996

**Authors:** Natalie G. Allen, Laahirie Edupuganti, David J. Edwards, Nicole R. Jimenez, Gregory A. Buck, Kimberly K. Jefferson, Jerome F. Strauss, Vaginal Microbiome Consortium, Edmond P. Wickham, Jennifer M. Fettweis

## Abstract

**Objective:** Evaluate differences between the vaginal microbiome of reproductive-aged women with overweight and obesity (Ow/Ob) compared with healthy weight (HW).

**Methods:** A cohort of 367 non-pregnant women (18-40 years) with Ow/Ob (body mass index [BMI] ≥25 kg/m^2^) was case-matched with 367 women with HW (BMI 18.0-24.9 kg/m^2^). The study was a secondary analysis of 16S rRNA vaginal microbiome surveys through the Vaginal Human Microbiome Study (VaHMP). Groups were matched on age, race/ethnicity, income, and nulliparity status.

**Results:** Mean age and BMI of Ow/Ob and HW groups were 26.8 versus 26.7 years and 37.0 versus 22.1 kg/m^2^, respectively. The overall vaginal microbiome composition differed between groups (PERMANOVA, p=0.035). Women with Ow/Ob had higher alpha-diversity compared with women with HW (Wilcoxon test, Shannon index p=0.025; Inverse Simpson index p=0.026). *Lactobacillus* dominance (≥30% proportional abundance) was observed in a greater proportion of women with HW (48.7%) compared with Ow/Ob (40.1%; p=0.026).

**Conclusions:** The vaginal microbiome differs in reproductive-aged women with Ow/Ob compared with women with HW, with increased alpha-diversity and decreased predominance of *Lactobacillus*. Observed differences in vaginal microbiome may partially explain differences in preterm birth and bacterial vaginosis risk between these populations.

## INTRODUCTION

There has been growing interest in elucidating the role of the microbiome in human health and disease. Mounting evidence implicates perturbations in the human microbiome as contributing factors to various disease states and conditions that affect both adults and children, including metabolic disorders such as overweight and obesity.^1,2^ The human microbiome is not homogenous, with the human body harboring distinct microbial populations across body sites, including the skin, oral, vaginal, and gut microbiomes.^3^ Although the health impacts of changes in the vaginal microbiome have been characterized less frequently than other body sites, the composition of the vaginal microbiome appears to have potentially important implications for a woman’s reproductive health.

The maternal microbiome also appears to have a significant impact on the health of offspring, resulting not only from the potential impact of the microbiome on maternal health, but also by its role in shaping the child’s developing microbiome. For example, variations in the maternal gut microbiome during pregnancy have been implicated in preterm birth and the development of allergies, asthma and obesity in offspring.^4-6^ Like the maternal gut microbiome, the vaginal microbiome is specifically thought to have implications for pregnancy outcomes including preterm birth;^7^ and, the composition of the maternal vaginal microbiome likely influences the child’s subsequent microbiome development after birth.^8-10^

Of note, differences in the composition of the vaginal microbiomes have been observed between women of African ancestry and European ancestry, with higher alpha-diversity in women of African ancestry and greater *Lactobacillus* dominance in women of European ancestry.^11,12^ As there have also been disparities noted in gynecological and pregnancy outcomes between these two racial groups, with higher rates of bacterial vaginosis (BV) and preterm birth noted twice as frequently in women of African ancestry,^13,14^ it is possible that race-associated differences in the composition of the vaginal microbiome contribute to the observed differences in incidence of adverse outcomes.

Obesity may also represent an additional factor that contributes to perturbations in the vaginal microbiome. Rates of obesity are higher in non-Hispanic Black women compared with non-Hispanic White women^15^ and among women with polycystic ovary syndrome (PCOS) compared with women without the condition.^16^ Moreover, women with obesity have higher rates of BV and adverse pregnancy outcomes, including preterm birth, compared with healthy-weight controls.^17-19^

In light of these observations, we endeavored to characterize whether alterations in the vaginal microbiome are associated with weight status in an effort to better understand the potential implications on pregnancy and neonatal outcome as well as overall reproductive health. To this end, we conducted a retrospective analysis of 16S rRNA sequencing data previously generated as part of the Vaginal Human Microbiome Project (VaHMP) to compare the vaginal microbiome of non-pregnant, reproductive-age women with overweight/obesity (Ow/Ob) and with healthy weight (HW).

## METHODS

### Vaginal Human Microbiome Project (VaHMP)

The Vaginal Human Microbiome Project at Virginia Commonwealth University was funded through the National Institutes of Health (NIH) as part of the NIH Human Microbiome Project (NIH grant UH3AI083263). In the VaHMP study, women were enrolled from obstetrics and gynecology clinics in Virginia as previously described under IRB no. HM12169 at Virginia Commonwealth University.^11^ Briefly, vaginal samples were collected with BD BBL CultureSwab EZ swabs (Becton, Dickerson and Company; Franklin Lakes, NJ) by rolling them on the vaginal sidewall about halfway between the introitus and the cervix, and processed for sampling within 4 hours of collection using the MO BIO PowerSoil DNA Isolation Kit (QIAGEN; Germantown, MD). DNA in the samples was amplified via PCR with primers targeted to the V1-V3 region of the 16S rRNA. The forward primer was a mixture (4: 1) of primers Fwd-P1 (5′-CCATCTCATCCCTGCGTGTCTCCGACTCAGBBBBBBAGAGTTYGATYMTGGCTYAG) and Fwd-P2 (5′-CCATCTCATCCCTGCGTGTCTCCGACTCAGBBBBBBAGARTTTGATCYTGGTTCAG and the reverse primer was Rev1B (5′-CCTATCCCCTGTGTGCCTTGGCAGTCTCAG. The PCR products were sequenced on the 454 GS FLX Titanium sequencer (Hoffmann-La Roche, LTD; Basel, Switzerland). Through the Research Alliance for Microbiome Science Registry (IRB no. HM15528), a subset of 16S rRNA data from the VaHMP study (IRB no. HM12169) was reanalyzed for the current analysis.

### Patient selection

Study participants with overweight or obesity (Ow/Ob), defined by body mass index (BMI) greater than 25 kg/m^2^, were matched with healthy weight (HW) controls, definedby BMI 18.0-24.9 kg/m^2^. Included participants were of reproductive age between 18 and 40 years (yr). Women in the Ow/Ob group were matched with HW counterparts based on age, race/ethnicity, socioeconomic status, highest level of education obtained, nulliparity status and clinic location using in-house Python scripts.

History was obtained by participant report in the VaHMP study via questionnaires that included race/ethnicity, yearly income bracket, educational level, gynecological history, sexual history, and medical history. Height and weight for BMI calculation were obtained from the clinical gynecology visit at the time that other data collection occurred.

### Bioinformatics and Statistical analyses

Reads that met the following criteria were processed: (1) valid primer and multiplex identifier sequences were observed; (2) less than 10% of base calls had a quality score less than 10; (3) the average quality score was greater than Q20; and (4) the read length was between 200 and 540 bases. Sequences were classified using STIRRUPS,^20^ an analysis platform that employs a global alignment algorithm combined with a curated vaginal 16S rRNA gene database.

Read counts of taxa classified at above threshold (97% identity) by STIRRUPS and present in at least 2% of the samples at > 0.1 % abundance were log transformed and normalized using the following formula:

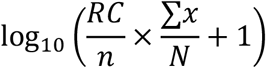

where RC represents the number of raw reads per taxon per sample, *n* is the number of sequences in a sample, the sum of *x* is the total number of counts of all samples and *N* is the total number of samples.^20^

After filtering out less abundant taxa using the above-mentioned steps, alpha-diversity measures (Shannon index and Inverse Simpson index) and PERMANOVA were calculated using diversity function and adonis function respectively via the R package vegan.^21^

Vagitype refers to the dominant taxon present in a sample with at least 30% proportional abundance. To assign samples to vagitypes, proportional abundance of taxa was used (not log transformed). The filtering step was the same as previously mentioned.

To explore taxa that contribute to differences in HW versus Ow/Ob groups, we made use of an L_1_ regularized logistic regression.^22^ That is, we sought a vector of weights, β, that minimizes 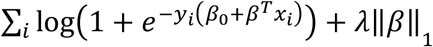, where y_*i*_ is the response (*i*.*e*., HW or Ow/Ob) and *x*_*i*_ are log-transformed taxa relative abundances. The constant *λ* is obtained via 10-fold cross validation. Taxa with non-zero weights were selected as potential contributors of group differences.

## RESULTS

A cohort 367 nonpregnant women of reproductive age (18-40 yr) with Ow/Ob was case-matched 1:1 with a cohort of 367 women with HW according to age, race/ethnicity, socioeconomic status, highest level of education and nulliparity status. Demographics of both groups and the overall cohort are described in Table 1. Of the 367 matched-pairs, 72.2% self-reported African ancestry. Mean age and body mass index (BMI) of the Ow/Ob and HW groups were 26.8 versus 26.7 yr, and 37.0 versus 22.1 kg/m^2^, respectively. The study population had a high prevalence of self-reported low annual income, with women in the Ow/Ob and HW reporting less than $20,000 yearly income in 61.4% and 58.3% of the cohort, respectively. Educational backgrounds are also reported, with 35.8% of the HW group and 29.7% of the Ow/Ob group noting completion of a college degree.

**Table 1.** Demographics of Healthy Weight (HW) and Overweight/Obesity (Ow/Ob) Groups. Race, ethnicity, income and educational status are based on self-report. BMI – body mass index. *HW and Ow/Ob groups were matched exactly on race and ethnicity.

| Demographic | All Participants<br>(n=734) | Healthy Weight<br>(n=367) | Overweight/Obesity<br>(n=367) | p-value (HW<br>vs Ow/Ob) |
| --- | --- | --- | --- | --- |
| Mean age, years<br>(standard deviation) | 26.8 (5.3) | 26.7 (5.3) | 26.8 (5.2) | 0.726 |
| Mean BMI, kg/m<sup>2</sup><br>(standard deviation) | 29.5 (8.6) | 22.1 (1.8) | 37 (5.8) | <0.001 |
| Race, n (%) |  |  |  |  |
| African ancestry | 530 (72.4) | 265 (72.4) | 265 (72.4) | 1.000* |
| European ancestry | 170 (23.2) | 85 (23.2) | 85 (23.2) | 1.000* |
| Other | 2 (0.3) | 1 (0.3) | 1 (0.3) | 1.000* |
| Hispanic, n (%) | 30 (4.1) | 15 (4.1) | 15 (4.1) | 1.000* |
| Annual income<br>bracket, n (%) |  |  |  |  |
| <\$20,000 | 414 (60) | 199 (58.4) | 215 (61.4) | 0.456 |
| \$20,000-40,000 | 127 (18.4) | 68 (20) | 59 (16.8) | 0.343 |
| \$40,000-60,000 | 73 (10.5) | 35 (10.3) | 38 (10.8) | 0.897 |
| \$60,000-80,000 | 32 (4.6) | 15 (4.4) | 17 (4.8) | 0.916 |
| >\$80,000 | 45 (6.5) | 24 (7) | 21 (6) | 0.691 |
| Education, n (%) |  |  |  |  |
| ≤High school | 333 (45.6) | 164 (45.2) | 169 (46) | 0.872 |
| Some college | 158 (21.6) | 69 (19) | 89 (24.3) | 0.103 |
| College graduate | 239 (32.8) | 130 (35.8) | 109 (29.7) | 0.093 |

Pertinent details of the gynecological histories of women in the HW and Ow/Ob groups are outlined in Table 2. The Ow/Ob and HW groups had a history of PCOS, BV, and smoking in 14.2% vs 9.5%, 37.8% vs 40.5%, and 53.9% vs 63% respectively.

**Table 2.** Gynecological History. Gynecological histories of all participants are reported according to healthy weight (HW) and overweight/obesity (Ow/Ob) groups including nulliparity status, contraceptive method, reported vaginal pH, history of PCOS and BV, history of smoking, number of sexual partners, and frequency of douching. Not progestin-only oral contraception (Oral Contraceptive [∅P]); Progestin-only oral contraception (Oral Contraceptive [P]); Progesterone-only injection (DMPA); Progesterone-only intrauterine device (LNG-IUS); Bacterial vaginosis (BV); Polycystic ovary syndrome (PCOS). * HW and Ow/Ob groups were matched exactly on nulliparity status.

|  | All Participants<br>(n=734) | Healthy Weight<br>(n=367) | Overweight/Obesity<br>(n=367) | p-value(HW<br>vs Ow/Ob) |
| --- | --- | --- | --- | --- |
| Nulliparity status, n (%) | 262 (35.7) | 131 (35.7) | 131 (35.7) | 1.000* |
| Contraceptive Method |  |  |  |  |
| Oral Contraceptive (ØP) | 110 (15.4) | 59 (16.5) | 51 (14.3) | 0.469 |
| Oral Contraceptive (P) | 5 (0.7) | 4 (1.1) | 1 (0.3) | 0.369 |
| DMPA | 94 (23.1) | 61 (29.3) | 33 (16.6) | 0.003 |
| LNG-IUS | 123 (30.2) | 46 (22.1) | 77 (38.7) | <0.01 |
| Non-hormonal Method | 252 (35.7) | 129 (36.5) | 123 (35) | 0.715 |
| PCOS, n (%) | 77 (12) | 31 (9.5) | 46 (14.2) | 0.086 |
| Current smoker, n (%) | 180 (58.4) | 97 (63) | 83 (53.9) | 0.133 |
| Vaginal pH, mean | 5.2 | 5.15 | 5.2 | 0.216 |
| Sexual partners (>1 in<br>past month), n (%) | 40 (6.2) | 21 (6.5) | 19 (6) | 0.927 |
| Douching (last month), n<br>(%) | 88 (13.5) | 42 (13) | 46 (14.1) | 0.791 |
| BV history, n (%) | 263 (38.3) | 136 (39.4) | 127 (37.1) | 0.591 |
| Current BV, n (%) | 159 (22) | 81 (22.1) | 78 (21.3) | 0.858 |

Contraceptive methods did differ significantly by weight status, with higher usage rates of progestin-only intrauterine devices (LNG-IUS) and lower rates of progestin-only injections (DMPA) in the Ow/Ob cohort. There were not significant differences noted in use of estrogen-containing contraceptive methods between groups, with 14.3% of the Ow/Ob cohort and 16.5% of the HW cohort using oral contraception that was not progestin-only.

The overall vaginal microbiome composition differed between the HW and Ow/Ob groups (PERMANOVA, p=0.035) after adjusting for other demographic and gynecological variables (Table 3). Specifically, *Lactobacillus* dominance differed between the two BMI groups (p=0.026) as shown in Figure 1. Among women with HW, 48.2% (177/367) of women exhibited a vaginal microbiome dominated by *Lactobacillus* species (defined at 30% or greater of total proportional abundance). In comparison, 40.1% (147/367) of women with Ow/Ob exhibited a vaginal microbiome dominated by a *Lactobacillus* species. Moreover, due to previously noted differences in vaginal microbiome composition based on race/ethnicity^11,12^ (Table 3), we conducted separate analyses stratified by race. In women of African ancestry, the difference in lactobacilli dominance according to weight status was significant (p=0.002). However, considering only women reporting European ancestry, a significant difference in lactobacilli dominance between the two weight categories was not observed (p=0.871). Of note, there was a higher incidence of lactobacilli dominance in the HW group in women of European ancestry (56/85; 66%) than women of African ancestry (114/265; 43%) as reported in previous studies.^11,12^

**Table 3.** PERMANOVA Table. Microbiome profiles are shown to be different among healthy weight (HW) and overweight/obesity (Ow/Ob) groups (p=0.035) when adjusting for other demographic and gynecological variables. Microbiome profiles also differ based on ethnicity/race (p=0.001), nulliparity status (p=0.001), and history of bacterial vaginosis (p=0.023).

| Source | Degrees of freedom | Sums of Squares | F ratio | p-value |
| --- | --- | --- | --- | --- |
| BMI Group | 1 | 0.563 | 2.39 | 0.035 |
| Ethnicity/Race | 3 | 2.223 | 3.14 | 0.001 |
| Annual Income | 5 | 1.610 | 1.37 | 0.093 |
| Age | 1 | 0.269 | 1.14 | 0.325 |
| Nulliparity Status | 1 | 2.081 | 8.82 | 0.001 |
| Contraception | 1 | 0.212 | 0.90 | 0.469 |
| PCOS Status | 2 | 0.415 | 0.88 | 0.566 |
| Sexual Partners | 1 | 0.328 | 1.39 | 0.227 |
| Douching | 1 | 0.159 | 0.67 | 0.673 |
| BV Status | 2 | 0.915 | 1.94 | 0.023 |
| Error | 478 | 112.693 |  |  |
| Total | 496 | 128.530 |  |  |

**Figure 1.**
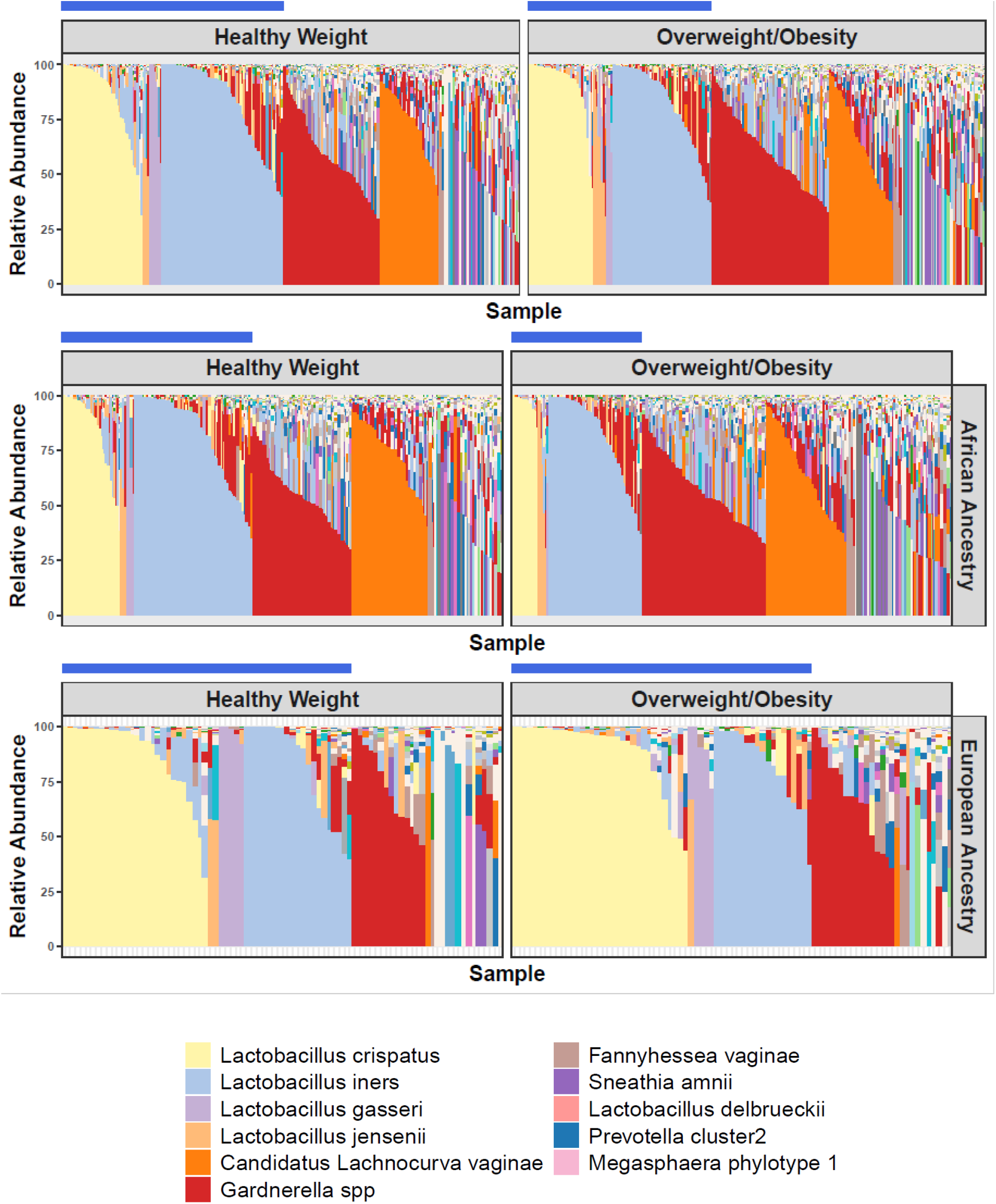
Vagitypes in Healthy Weight (HW) versus Overweight/Obesity (Ow/Ob) by Race. Vagitypes of women in the healthy weight and overweight/obesity groups according to total cohort, African ancestry, and European ancestry. The blue bar identifies vagitypes with *Lactobacillus* dominance (at least 30% proportional abundance).

As shown in Figure 2, women with Ow/Ob also had higher vaginal microbiome alpha-diversity compared with women with HW, as measured by either Shannon index (p=0.025) or Inverse Simpson index (p=0.026). When stratified into women of African ancestry and European ancestry, the increase in alpha-diversity was significant in Ow/Ob women of African ancestry (p=0.025, Shannon Index; p=0.025, Inverse Simpson), but not in women of European ancestry (p=0.15, Shannon Index; p=0.15, Inverse Simpson). When subdivided into nulliparous versus women who had previously given birth, there was no significant difference in alpha diversity by both Shannon index and Inverse Simpson index (data not shown).

**Figure 2.**
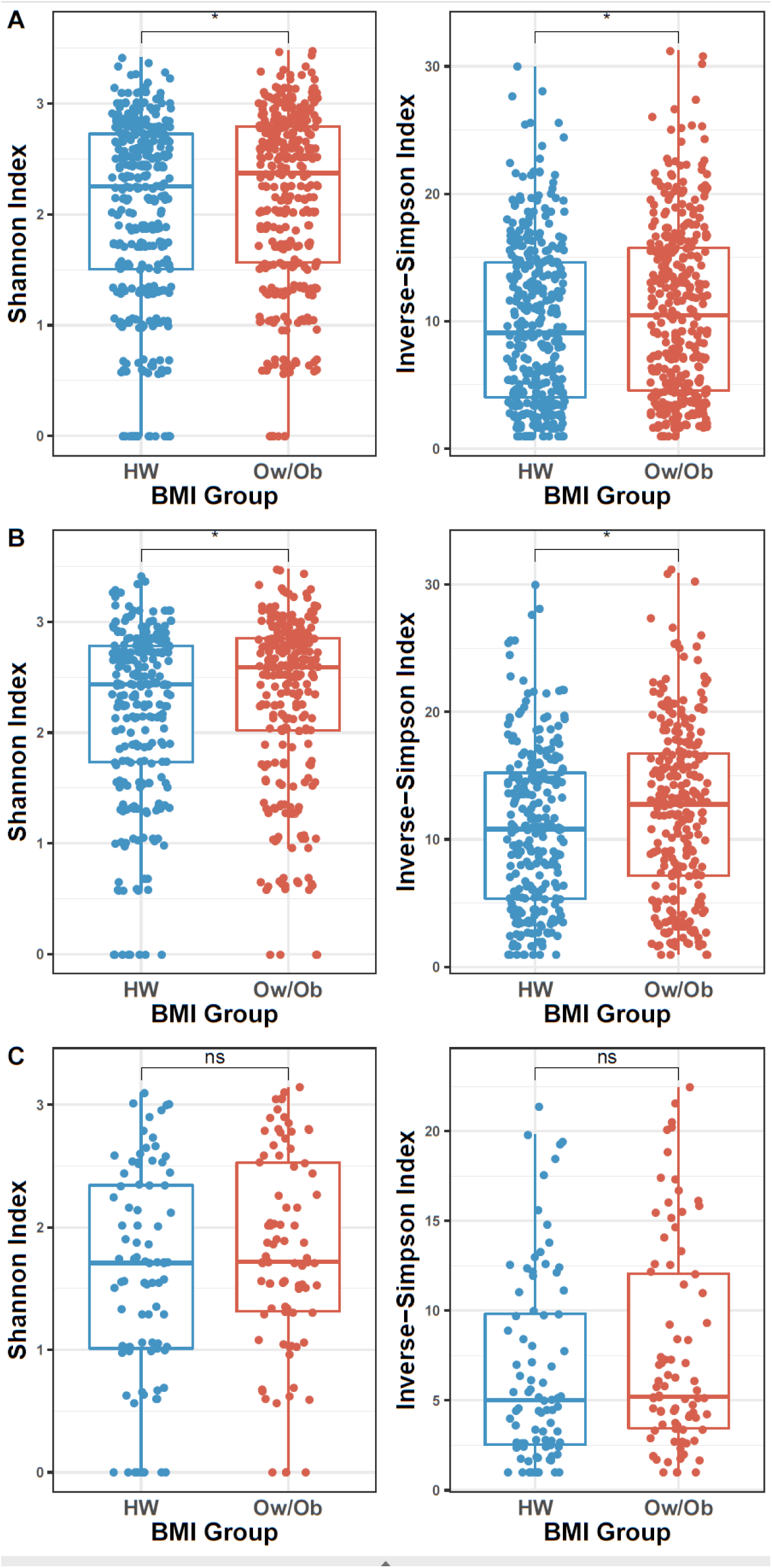
Vaginal Microbiome Diversity Measures in Women with Healthy Weight (HW) versus Overweight/Obesity (Ow/Ob) by Race. A. Shannon index and Inverse-Simpson Index were significantly different in HW (blue) compared with Ow/Ob (red) considering the entire cohort; B. Among those of African ancestry, alpha-diversity was higher in the Ow/Ob group than in HW; C. In participants of European ancestry, there was no significant difference in alpha-diversity between HW and Ow/Ob groups.

The regularized logistic regression model revealed multiple taxa of interest that contribute to differences between the HW cohort and the Ow/Ob cohort. Taxa identified as contributing to these differences were Coriobacteriaceae OTU27, *Megasphaera* OTU71_type1 (recently renamed Megasphaera lornae sp. nov),^23^ Saccharibacteria TM7 OTUH1, Prevotellaceae OTU61, and *Mobiluncus mulieris* (Figure 3). The full heat map analysis is noted in Figure S1.

**Figure 3.**
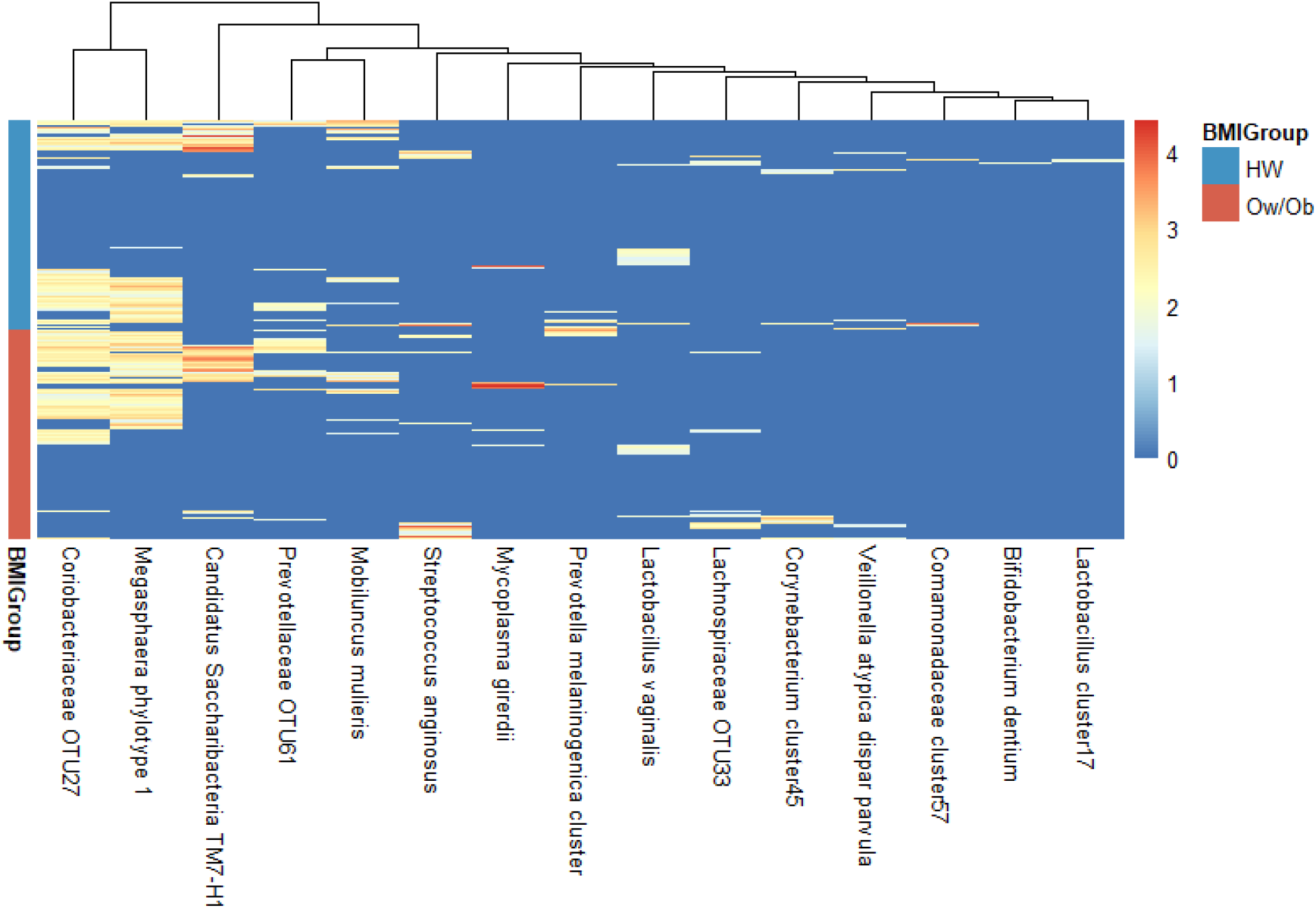
Heatmap of Taxa Identified in Regularized Logistic Regression Model. Taxa identified by regularized logistic regression model to have nonzero weights and to be potential contributors of differences between healthy weight (HW) and overweight/obesity (Ow/Ob) cohorts.

## DISCUSSION

The vaginal microbiome composition, including *Lactobacillus* dominance, alpha-diversity, and specific taxa, differs between women of reproductive age with overweight and obesity (Ow/Ob) compared with women with healthy weight (HW). In addition to weight status, race/ethnicity, nulliparity status, and history of BV were associated with significant differences in the vaginal microbiome. The causes of the observed differences due to weight status and any implications for subsequent gynecological and pregnancy outcomes are still unclear; however, there are likely contributions from genetic background and other environmental factors. Within the study, the matching process controlled for the potential impact of race/ethnicity, income, and previous birth history, all factors previously known to influence the vaginal microbiome.^24^

In our study, women with Ow/Ob exhibited less lactobacilli dominance than women with HW. Prototypical health of the vaginal microbiome has been thought to consist of a vaginal profile that is dominated by lactobacilli and a low pH, factors that inhibit the growth of other bacteria. When perturbations occur, dysbiotic states may arise like BV, the most common vaginal condition in women of childbearing age.^25^ In addition, BV has also been associated with increased rates of sexually transmitted infections and pregnancy complications including preterm birth.^26-28^

We also observed greater alpha-diversity among women with Ow/Ob compared with women with HW. Alpha-diversity typically decreases in pregnancy, and an increase in alpha-diversity prior to pregnancy has previously been associated with adverse pregnancy outcomes including preterm birth.^29^

Specific taxa were identified to be more prevalent in the Ow/Ob cohort including Coriobacteriaceae *OTU27, Megasphaera* OTU71_type1, Saccharibacteria TM7 OTUH1, Prevotellaceae OTU61, and *Mobiluncus mulieris*. Both *Megasphaera OTU71_type1* and Coriobacteriaceae, although not specifically OTU27, have been implicated in BV.^30^ Several of the taxa identified more frequently in the Ow/Ob group have also been recently associated with preterm birth including *Megasphaera* OTU71_type1, Saccharibacteria TM7 OTUH1, and *Mobiluncus mulieris*.^31,32^ Also, in a preterm birth study, TM7 OTUH1 was shown to have a positive association with Coriobacteriaceae OTU27, Megasphaera OTU71_type1, and Prevotellaceae OTU61.^32^ Thus, this group of bacteria may co-associate with one another in certain conditions. Differences noted in the vaginal microbiome between the HW and Ow/Ob groups may help to shed light on a component of the increased risk in women with obesity for adverse gynecological and obstetrical outcomes.^33^

In addition to weight status, our analyses confirmed that race/ethnicity was a significant predictor of differences in the vaginal microbiome. As a result, we conducted secondary analyses of the impact of race/ethnicity in addition to the overall group. Alpha-diversity was noted to be higher in women in the Ow/Ob group than in women in the HW group, both in the overall group and in the subset of women of African ancestry. Increased alpha-diversity has been previously reported in women of African ancestry compared with European ancestry.^11^ Of note, increased alpha-diversity has also been associated with increased rates of BV and preterm birth in some studies.^29,34^

We also observed differences in *Lactobacillus* dominance in the vaginal microbiome profiles between women in the Ow/Ob and HW groups, in both the overall cohort and among women of African ancestry. Differences in the prevalence of lactobacilli dominance of the vaginal microbiome between women of African versus European ancestry with HW have been previously described.^11^ These differences in lactobacilli dominance between women in Ow/Ob versus HW may have significant implications for gynecological health.

The composition of vaginal microbiome is influenced by many factors including systemic estrogen levels.^35^ Estrogen induces maturation of the vaginal epithelium and increases in glycogen deposition^35^ and is thought to drive significant changes in the vaginal microbiome across different life stages. For many women, *Lactobacillus* dominance persists through their reproductive years and concordantly decreases as estrogen levels decline during menopause.^36^ With estrogen likely representing a major mediator on the vaginal microbiome, scenarios that lead to altered estrogen levels may further impact the composition of the vaginal microbiome. Women using estrogen-containing combined oral contraceptives (COC) have been noted to have a greater abundance of *Lactobacillus* compared with women who used DMPA or LNG-IUS.^37,38^ Studies have also demonstrated increased *Lactobacillus* in post-menopausal women taking hormone replacement therapy (HRT) as opposed to those without HRT.^39,40^

In addition to the pharmacologic influence of hormonal contraception or treatment of the vaginal microbiome, researchers have identified changes in the vaginal microbiome in association with the high-estrogen state of pregnancy, specifically noting a decreased diversity of species and enhanced *Lactobacillus* dominance^11^ that has been shown to persist at least one year following delivery.^24^ Other sex steroids may also impact vaginal microbiome composition. PCOS, a common endocrine disorder among women of reproductive age, is characterized by irregular menstrual cycles and hyperandrogenemia. A recent study conducted by Hong and colleagues investigated the association between PCOS status and the vaginal microbiome, and noted a lower relative abundance of *Lactobacillus* in PCOS samples when compared with the control group, after controlling for BMI and androgen levels.^41^

Obesity has been associated with elevated estrogen levels across both sexes,^42,43^ resulting from enhanced peripheral aromatization of androgens to estrogens in adipose tissue. Thus, our observation that women with Ow/Ob have less *Lactobacillus* dominance may seem counter-intuitive. However, in reproductive age women, such as our study population, the aromatization resulting from obesity appears to have negligible effect on systemic estrogen levels.^44^ Thus, the impact of obesity on the vaginal microbiome in women of reproductive age is likely independent of estrogen effect. In addition to the hormonal status, the presence of other factors including nulliparity, tobacco use, socioeconomic status and race/ethnicity have been associated with differences in the vaginal microbiome.^24,45^

The current study design had some consistent strengths, including relatively large cohorts of reproductive age women with Ow/Ob and HW. Additionally, the exclusion criteria allowed us to separate possible effects from type 1 and type 2 diabetes, gestational diabetes, and hysterectomy. The strength of the VaHMP cohort and previously performed 16S rRNA analysis permitted us to examine a large cohort in a timely manner. The study does have some limitations. Firstly, some medical history and demographic information was by self-report, allowing for possible recall or classification bias from participants. Another limitation is that not all races/ethnicities were well represented in the study population. Moreover, of available participants, smaller numbers of women reported European ancestry which may have impacted the ability to detect significant differences in the vaginal microbiome in this specific group.

## CONCLUSIONS

Our findings contribute to the literature in the vaginal microbiome field by demonstrating that vaginal microbiome composition differs between women with healthy weight (HW) versus overweight and obesity (Ow/Ob) in multiple measures, including *Lactobacillus* dominance, alpha-diversity, and specific taxa. Women in the Ow/Ob group, had lower rates of *Lactobacillus* dominance and lower alpha diversity, both of which are linked to adverse gynecological and pregnancy outcomes. Some specific taxa noted to be increased in Ow/Ob have been implicated in increased rates of preterm birth. Differences noted in the vaginal microbiome between the HW and Ow/Ob groups may help to shed light on a component of the increased risk in women with obesity for adverse gynecological and obstetrical outcomes. Further studies are needed to determine whether the vaginal microbiome differences reported here are reproduced in other cohorts of women and if such changes modulate risk of gynecological and pregnancy outcomes among these groups.

## AUTHOR CONTRIBUTIONS

N.A. conducted all experiments and analyses and drafted the manuscript. L.E. performed case-matching and bioinformatic analyses. D. E. performed statistical analyses. N.R.J. assisted in manuscript preparation and clinical association analyses. G.A.B., K.K.J, J.F.S.III, and J.M.F. comprised the executive committee of the parent study. J.M.F. and E.P.W. supervised this study and led the overall direction and planning. N.A., E.P.W. and J.M.F. designed the study and wrote the manuscript with contributions from all other authors.

## ACKNOWLEDGMENTS

The study team gratefully acknowledges the participants who contributed specimens and data to the Vaginal Human Microbiome Project (VaHMP). The authors would also like to acknowledge other members of the Vaginal Microbiome Consortium and the Research Alliance for Microbiome Science (RAMS) Registry whose contributions made the study possible including the team of research coordinators, the team of sample processors, the team of data managers and the team of clinicians and nurses who assisted with sample collection. This study was funded by NIH grants R21HD092965 and U54HD080784. Other grants that provided partial support include NIH grant UH3AI083263 and a GAPPS BMGF PPB grant. NRJ was supported by NIH grant R25GM090084 for the VCU Initiative for Maximizing Student Development (IMSD) program. DNA sequencing was performed in the Genomics Core of Nucleic Acids Research Facilities at VCU; the Center for High Performance Computing at VCU provided supercomputing clusters, large-scale storage systems and support for high-performance computing.

## COMPETING INTERESTS

All authors declare no financial conflicts of interest.

---

**Figure S1:**
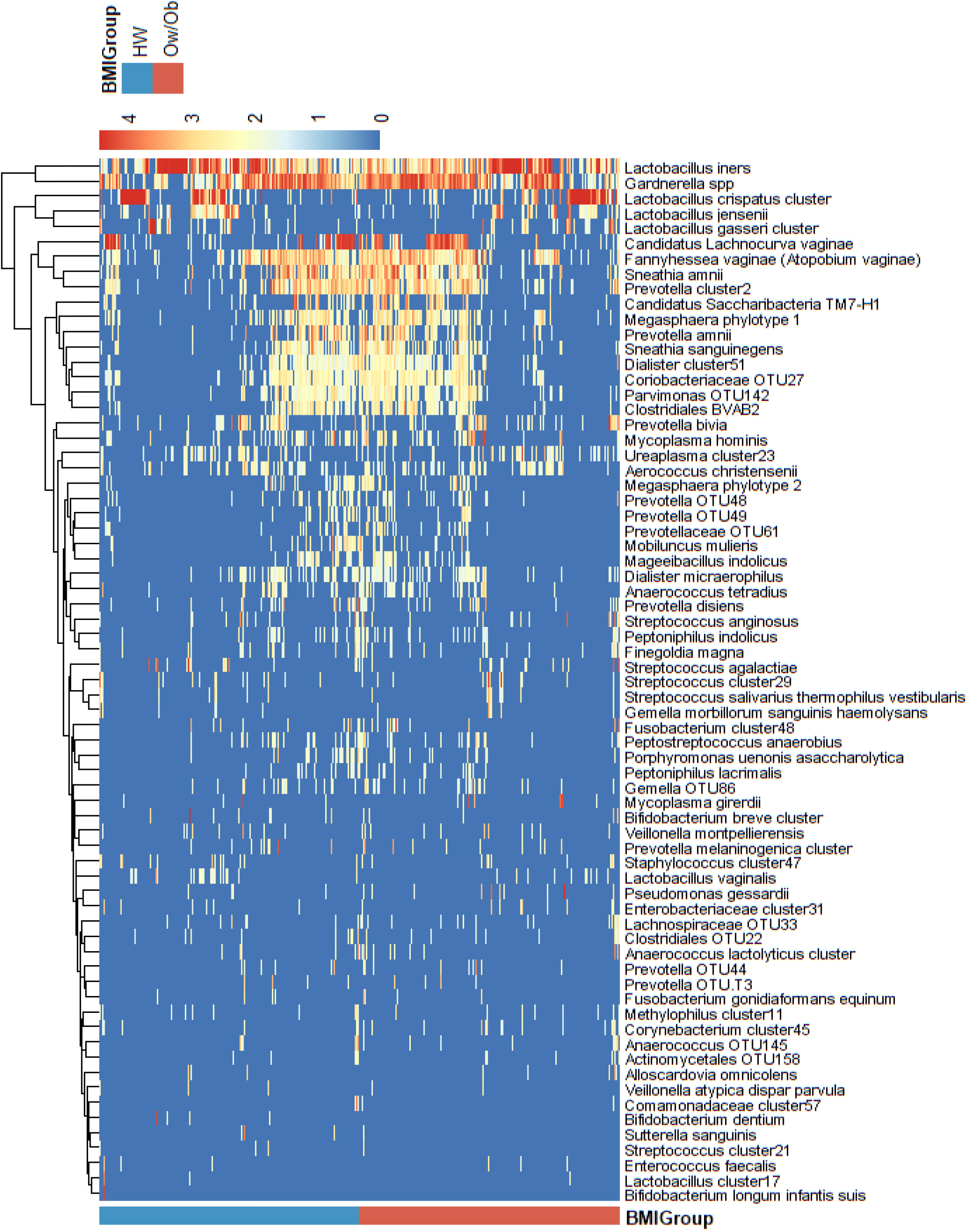
Heatmap of Taxa Identified by Weight Group. Taxa identified by regularized logistic regression model in the analysis between healthy weight (HW) and overweight/obesity (Ow/Ob) cohorts.

## Notes

### Competing Interest Statement

The authors have declared no competing interest.

